# Stimulated human cortical waves: spatiotemporal dynamics across development

**DOI:** 10.64898/2026.01.12.699175

**Authors:** Xi Wang, Bingjiang Lyu, Lang Qin, Jingjing Li, Yikun Zhang, Andrew Zalesky, Chunfeng Yang, Yang Chen, Miao Cao

## Abstract

Traveling waves of neural activity are fundamental to cortical function, yet their spatiotemporal dynamics and structural constraints remain less understood in human brain. Conventional analyses focus on network topology and spatial geometry of neural signals, with limited emphasis on wave dynamics at high spatiotemporal resolutions. Here, we applied a manifold-based optical flow algorithm to an open-access intracranial electroencephalography (iEEG) dataset from the single-pulse stimulation cortico-cortical evoked potentials (CCEPs) paradigm, enabling direct construction of high-temporal-resolution cortical neural activity velocity fields. Spatiotemporal mode decomposition revealed a hierarchical organization: large-scale plane waves (mediating global propagation) dominate, complemented by local patterns for local computation. Dominant plane wave directions correlate with underlying white matter fibre tract orientations, indicating anatomical guidance of cortical dynamics. Velocity field singular points (sources, sinks, spirals and saddles) are non-randomly distributed, associating with default mode network (DMN) hubs. We further identified a nonlinear relationship between cortical wave speed and age, with developmental phases paralleling white matter maturation and aging. Our findings link cortical wave dynamics to anatomy, functional networks, and neurodevelopment, establishing wave speed as a potential yet important biomarker for brain function and health.

**Significance Statement:** Traveling waves may coordinate neural activity across the cortex in a structured fashion, yet their organization remains less understood. Applying a manifold-based method to intracranial electroencephalography (iEEG) recordings across development, we demonstrate cortical waves, evoked by direct cortical stimulations, exhibit hierarchical spatiotemporal patterns, including dominant large-scale waves that mediate global communication and localized patterns that support regional computation. Wave propagation directions, correlating with underlying white matter fiber tracts, may also reveal structural foundations of cortical dynamics. More crucially, cortical wave speed varies nonlinearly across lifespan, associating with white matter development and aging. These findings may establish wave speed, a physical metric, as a promising biomarker linking brain anatomy, functional networks, and neurodevelopment, with implications for understanding neurological and psychiatric disorders.

## Introduction

How the intricate anatomical architecture of the brain gives rise to its dynamic functional organisation remains a central question in neuroscience. At the macro-scale, white-matter pathways constrain communication between distributed cortical regions, and structural connectivity can predict aspects of functional connectivity (***Honey et al., 2009; Suárez et al., 2020***). Yet structure and function are far from isomorphic, as functional interactions can arise through polysynaptic and network-mediated pathways (***Suárez et al., 2020***), while structure-function coupling varies markedly across the cortex and weakens along the cortical hierarchy towards transmodal regions (***Valk et al., 2022; Fotiadis et al., 2024***). This imperfect correlation suggests that anatomy constrains, rather than uniquely determines, the repertoire of functional dynamics emerging from nonlinear and recurrent network interactions (***Yang et al., 2023; Fotiadis et al., 2024***). How this balance between anatomical constraints and network dynamics is expressed when cortical activity is externally perturbed remains unclear. Direct cortical stimulation provides a unique opportunity to investigate this question by initiating activity at a defined cortical site, disturbing surrounding cortical activity and tracking its propagation through the connected network (***Keller et al., 2014; Lyu et al., 2025***). Unlike spontaneous activity, which emerges from ongoing distributed dynamics, stimulation imposes a temporally localised perturbation that may expose anatomical constraints otherwise obscured by endogenous activity. Previous observations reveal a fundamental question: does stimulation expose the physical constraints imposed by white-matter architecture, or does it reveal an emergent propagation regime in which anatomical constraints are transformed by recurrent, polysynaptic and hierarchical network dynamics? Would direct stimulation give us a unique window to speculate the wiring of white-matter structure and how cortical coordination is constrained by white-matter dynamically?

Cortical travelling waves offer a potential mechanistic link between structure and function (***Koller et al., 2024***). Unlike functional connectivity, which largely describes statistical dependencies between regions, travelling waves explicitly encode the spatiotemporal trajectory, direction and velocity of neural activity (***Muller et al., 2018; Zhang et al., 2018***). Waves have been observed across sensory, motor and cognitive systems and are proposed to coordinate neural computation across spatial scales (***Muller et al., 2018; Rubino et al., 2006; Zhang et al., 2018***). At the macro-scale, the propagation of rhythmic activity can reflect long-range conduction through white-matter pathways: human alpha waves exhibit structured cortical propagation (***Zhang et al., 2018***), and their dynamics have been related to anatomical connectivity and conduction delays (***Muller et al., 2016; Koller et al., 2024***). Beyond large-scale propagation, cortical waves can also exhibit complex local patterns, including sources, sinks and spirals (***Roberts et al., 2019; Das et al., 2026***). Thus, cortical waves provide a physical description of how anatomical connections may be transformed into time-varying patterns of communication.

In the present study, we aim to address critical questions in brain structure-function relationship by characterizing cortical activity as a spatiotemporal velocity field derived from human intracranial EEG (iEEG) recordings during cortico-cortical evoked potentials (CCEPs) (***Qiang et al., 2025***). We harness a manifold-based optical-flow framework that reconstructs the instantaneous velocity of cortical activity, enabling the spatial organization of propagation to be quantified beyond conventional amplitude- or connectivity-based measures. Here, we hypothesise that cortical propagation is hierarchically organised, with dominant large-scale modes supporting global com-munication and secondary localized patterns contributing to regional computation; that propagation dynamics are constrained by the underlying anatomical architecture, such that dominant wave directions tend to align with white-matter fibre-tract orientations; and that local dynamical singular points, including sources, sinks, spirals and saddles, provide sites at which local activity can reshape large-scale propagation. Finally, because white-matter architecture undergoes substantial maturation and ageing across the lifespan, we hypothesise that cortical wave speed follows a nonlinear developmental trajectory that parallels changes in white-matter organisation. By linking propagation direction, local dynamical structure and wave speed to brain anatomy across development, this approach seeks to establish a direct relationship between the structural architecture of the cortex and its emergent spatiotemporal dynamics.

## Results

Of the 74 patients in the CCEP dataset, 67 were retained after exclusions related to surface reconstruction or temporal segmentation (Methods and Supplementary Material S2.1). We applied a manifold-based optical flow method to their CCEP recordings to reconstruct stimulus-evoked cortical velocity fields (***Figure 1***). Electrode potentials were interpolated onto surface manifolds reconstructed from electrode coordinates, and successive potential fields were used to estimate local propagation velocities. We then characterized singular points and their spatial association with default mode network (DMN) regions. We also decomposed the velocity fields into spatiotemporal modes and compared the local velocity directions of the dominant spatial mode with projected tract connection axes. Finally, we examined the relationship between cortical wave speed and age.

**Figure 1.**
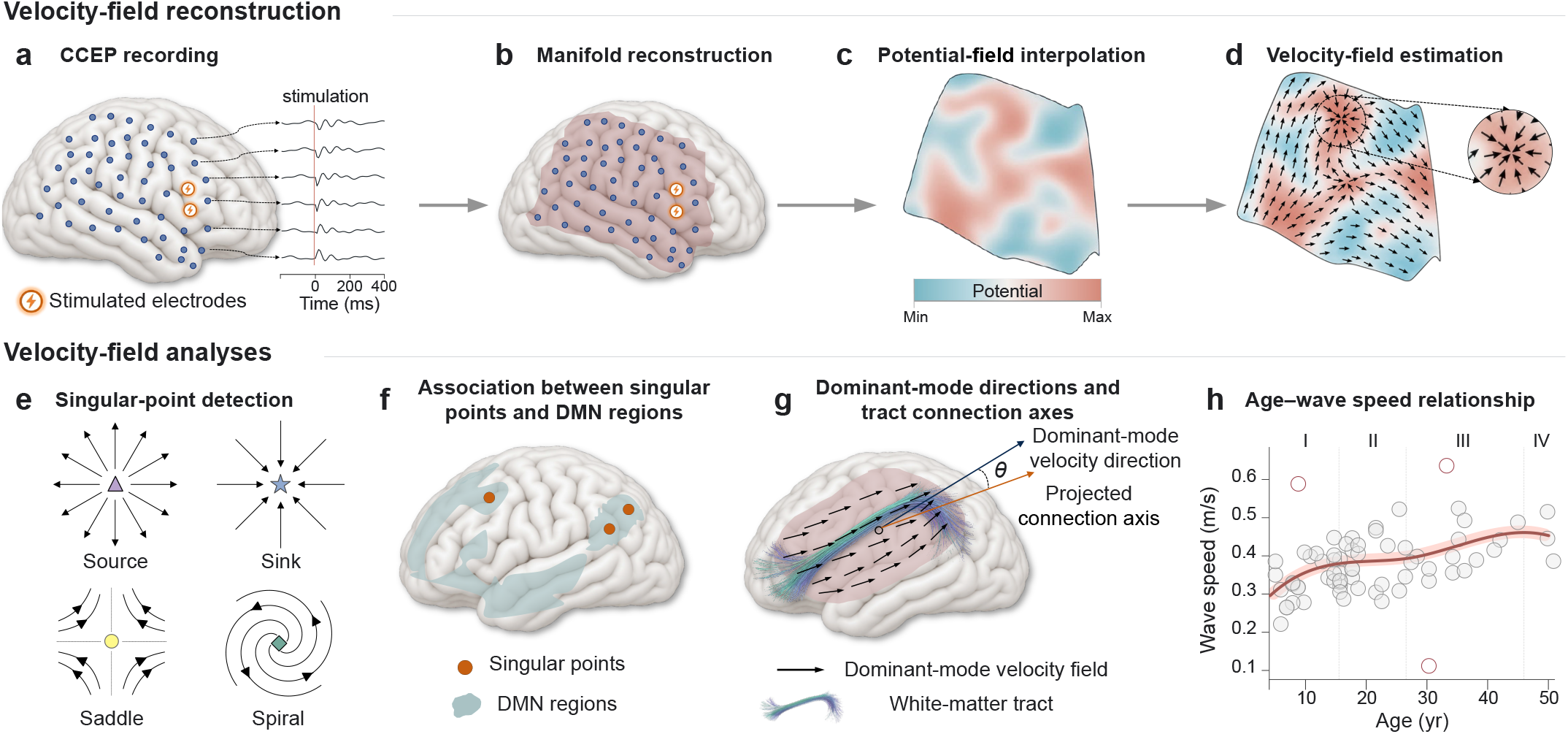
Workflow for velocity-field reconstruction and analysis of stimulus-evoked cortical waves. **a**, Single-pulse electrical stimulation is delivered through a pair of cortical electrodes, and cortico-cortical evoked potentials (CCEPs) are recorded from the remaining electrodes. Orange lightning symbols mark stimulated electrodes. **b**, A cortical surface manifold is reconstructed from electrode coordinates. **c**, Electrode potentials are interpolated onto the reconstructed manifold to generate continuous potential fields. Colors indicate potential amplitude. **d**, Manifold-based optical flow estimates velocity fields from successive potential fields. Black arrows indicate local velocity vectors; the inset highlights flow around a singular point. **e**, Schematic wave patterns illustrate four singular-point types: sources, sinks, saddles and spirals. **f**, The spatial association between detected singular points and default mode network (DMN) regions is examined. **g**, Directional correspondence between the dominant spatial mode (Mode 1) and the projected connection axes of long-range white-matter tracts. Each connection axis links the tract’s two cortical termination regions, and *θ* denotes the angle between its local tangent-plane projection and a Mode-1 velocity vector. **h**, Relationship between cortical wave speed and age. Points represent subject-level measurements, the solid curve shows the nonlinear fit and Roman numerals I–IV indicate four age intervals identified from the fitted curve.

### Cortical wave singular points and their spatial association with the DMN

We characterized local wave patterns by identifying singular points, where the reconstructed velocity field vanishes (***Perry and Chong, 1987***). Singular points were classified as sources, sinks, saddles or spirals according to the local geometry of the surrounding velocity field (Methods; ***Figure 1***e) (***Takens, 1974; Townsend and Gong, 2018***). In the patient recordings, representative source and saddle configurations were evident in the reconstructed velocity fields (***Figure 2***a). Examples included a source within a DMN region in Subject 01 and a saddle outside the DMN in Subject 02 (***Figure 2***b).

**Figure 2.**
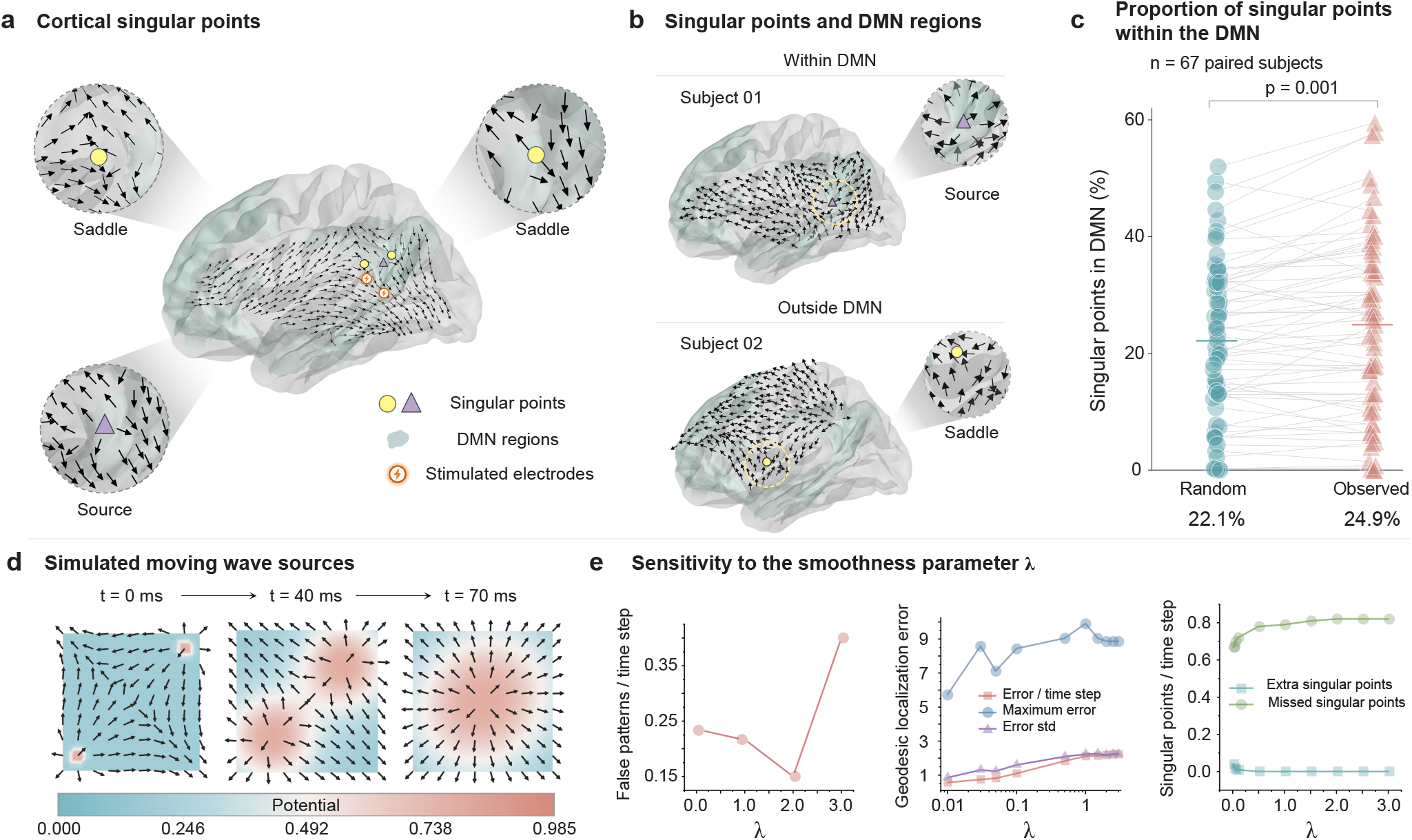
Detection of cortical wave singular points and their spatial association with the default mode network. **a**, Representative singular points detected from stimulus-evoked cortical velocity fields. Black arrows indicate local velocity vectors; purple triangles and yellow circles mark sources and saddles, respectively. Enlarged views show the surrounding wave patterns. Light-blue shading denotes default mode network (DMN) regions, and orange lightning symbols mark stimulated electrodes. **b**, Examples of a source within the DMN (Subject 01) and a saddle outside the DMN (Subject 02). **c**, Paired subject-level proportions of singular points within the DMN for observed data (coral triangles) and a within-manifold random baseline (cyan circles; *n* = 67 subjects). DMN regions were defined using the Yeo seven-network atlas. Gray lines connect paired values, and horizontal bars indicate cohort medians (random, 22.1%; observed, 24.9%). Observed proportions exceeded the random baseline (*p* = 0.001, one-sided Wilcoxon signed-rank test). **d**, Two simulated moving wave sources at 0, 40 and 70 ms. Colors indicate potential, and arrows show the corresponding velocity field. **e**, Sensitivity of singular-point detection to the velocity-field smoothness parameter *λ* in simulations with known ground truth. Left, false patterns per time step; middle, geodesic localization error (error per time step, maximum error and error standard deviation); right, extra and missed singular points per time step.

To assess detection performance, we analyzed synthetic data with known singular-point locations and types. Simulations included two moving Gaussian wave sources converging along diagonal trajectories (***Figure 2***d), together with stationary node, focus and saddle patterns (20 of each type; Methods). Varying the velocity-field smoothness parameter *λ* revealed different responses across detection metrics (***Figure 2***e). Increasing *λ* reduced extra singular-point detections but increased missed detections and generally increased geodesic localization error. False-pattern counts varied non-monotonically with *λ*, decreasing over part of the tested range before increasing at the largest value. These results illustrate a trade-off between suppressing extra detections and preserving localization accuracy and detection sensitivity. We selected *λ* = 0.01 for velocity-field estimation from the CCEP recordings, prioritizing localization accuracy and detection sensitivity (Methods).

We next examined whether singular points were enriched within DMN regions defined by the Yeo seven-network atlas (***Yeo et al., 2011***). For each subject, the observed proportion of singular points within the DMN was compared with a random baseline generated within the same reconstructed cortical manifold. Across 67 subjects, the cohort median was 24.9% for observed singular points and 22.1% for the random baseline (***Figure 2***c). Observed proportions were higher than their paired baseline values (*P* = 0.001, one-sided Wilcoxon signed-rank test). Paired results and additional methodological details are provided in Supplementary Material S2.5.

### Spatiotemporal modes of stimulus-evoked cortical velocity fields

While singular-point analysis characterizes local wave features, it does not quantify how different propagation patterns contribute to the overall stimulus-evoked dynamics. We therefore decomposed the reconstructed velocity fields into spatiotemporal modes, each comprising a spatial propagation pattern and a corresponding temporal profile (Methods). The relative contribution of each mode was quantified by its explained variance fraction. The spatial component associated with the largest fraction was termed the dominant spatial mode (Mode 1).

Across 67 subjects, the dominant spatial mode explained a mean of 72.97% of the variance (median, 73.51%; Supplementary Information, Section S2.2.4 and Table S7). In the representative recording from Subject 01, this mode explained 85.96% of the variance and exhibited broad, plane-wave-like propagation (***Figure 3***a). Secondary modes (Modes 2–4) accounted for 6.26%, 2.63% and 1.56%, respectively, and displayed more heterogeneous spatial patterns. The temporal profiles showed transient post-stimulus fluctuations that diminished toward the end of the analysis interval. The dominant spatial mode explained 88.89% and 85.14% of the variance in the representative recordings from Subjects 35 and 48, respectively (***Figure 3***b). These examples showed a common hierarchical organization: the dominant spatial mode exhibited broad, plane-wave-like propagation, whereas secondary modes captured more localized patterns, some containing sources, sinks, saddles or spirals. Propagation directions, singular-point locations and mode-specific variance fractions nevertheless differed among subjects. Given its large contribution to the reconstructed dynamics, we used the dominant spatial mode for the subsequent comparison with tract connection axes.

**Figure 3.**
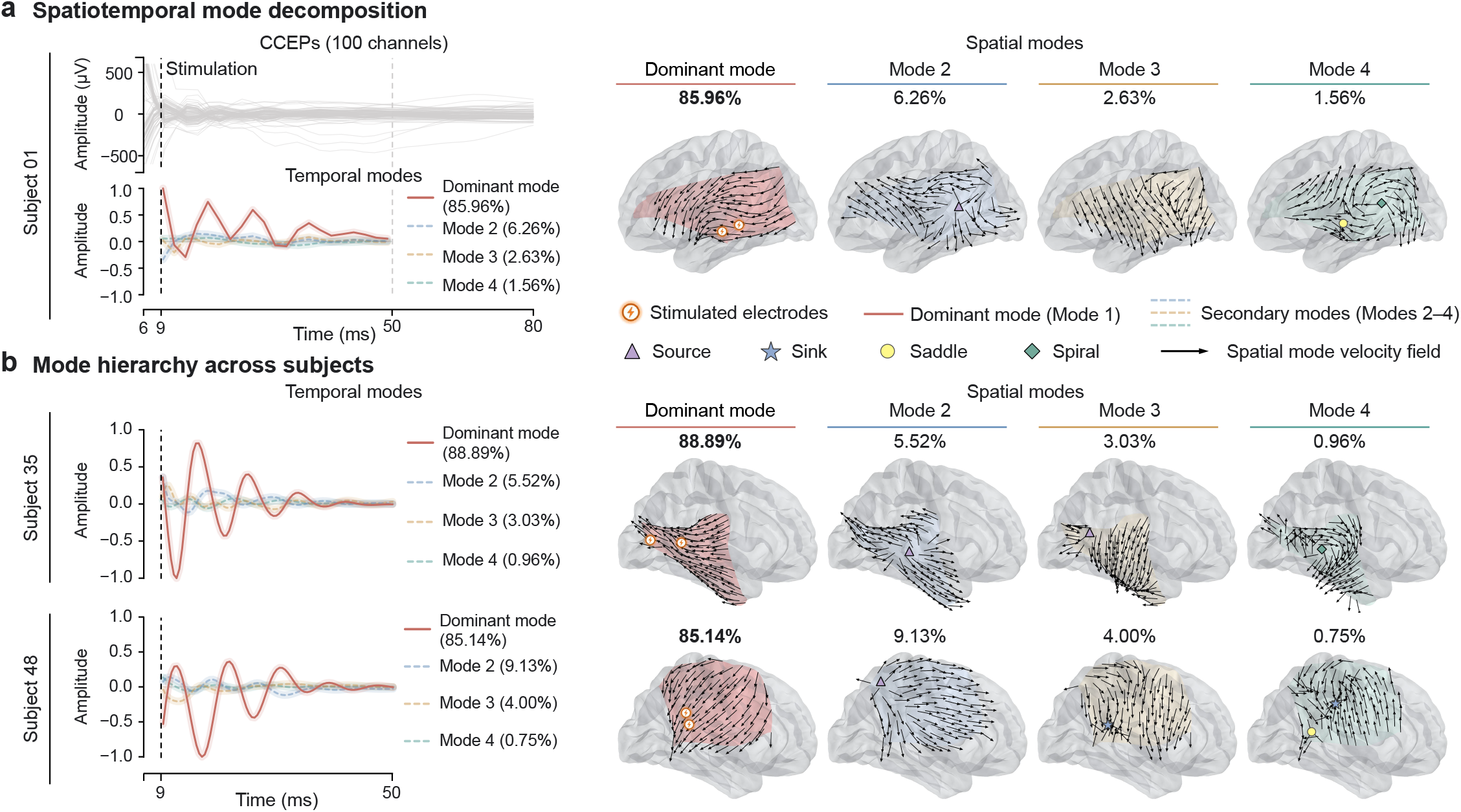
Hierarchical spatiotemporal modes of stimulus-evoked cortical wave propagation. **a**, Spatiotemporal mode decomposition of the reconstructed velocity fields for a representative recording from Subject 01. Cortico-cortical evoked potentials (CCEPs) from 100 recording channels (top left) are shown alongside the first four temporal modes (bottom left) and corresponding spatial modes (right). The dominant mode (Mode 1) explains 85.96% of the variance and exhibits large-scale, plane-wave-like propagation, whereas secondary modes (Modes 2–4) explain 6.26%, 2.63% and 1.56%, respectively, and capture more heterogeneous spatial patterns. **b**, Corresponding temporal modes (left) and spatial modes (right) for representative recordings from Subjects 35 (top) and 48 (bottom). The dominant mode explains 88.89% and 85.14% of the variance, respectively. Percentages indicate the variance explained by each mode in the illustrated recording. Black arrows indicate spatial-mode velocity vectors; purple triangles, blue stars, yellow circles and green diamonds mark sources, sinks, saddles and spirals, respectively. Orange lightning symbols mark stimulated electrodes.

### Association between the dominant spatial mode and tract connection axes

We next examined directional correspondence between the dominant spatial mode and the end-to-end orientation of long-range white-matter connections. The analysis included 65 subjects: 10 for the arcuate fasciculus (AF), 23 for the frontal–central segment of the superior longitudinal fasciculus (SLF), 9 for the frontal–parietal SLF and 23 for the temporo-parietal aslant tract (TPAT). Representative dominant-mode velocity fields are shown alongside these tracts for responses evoked by stimulation of different electrode pairs (***Figure 4***a).

**Figure 4.**
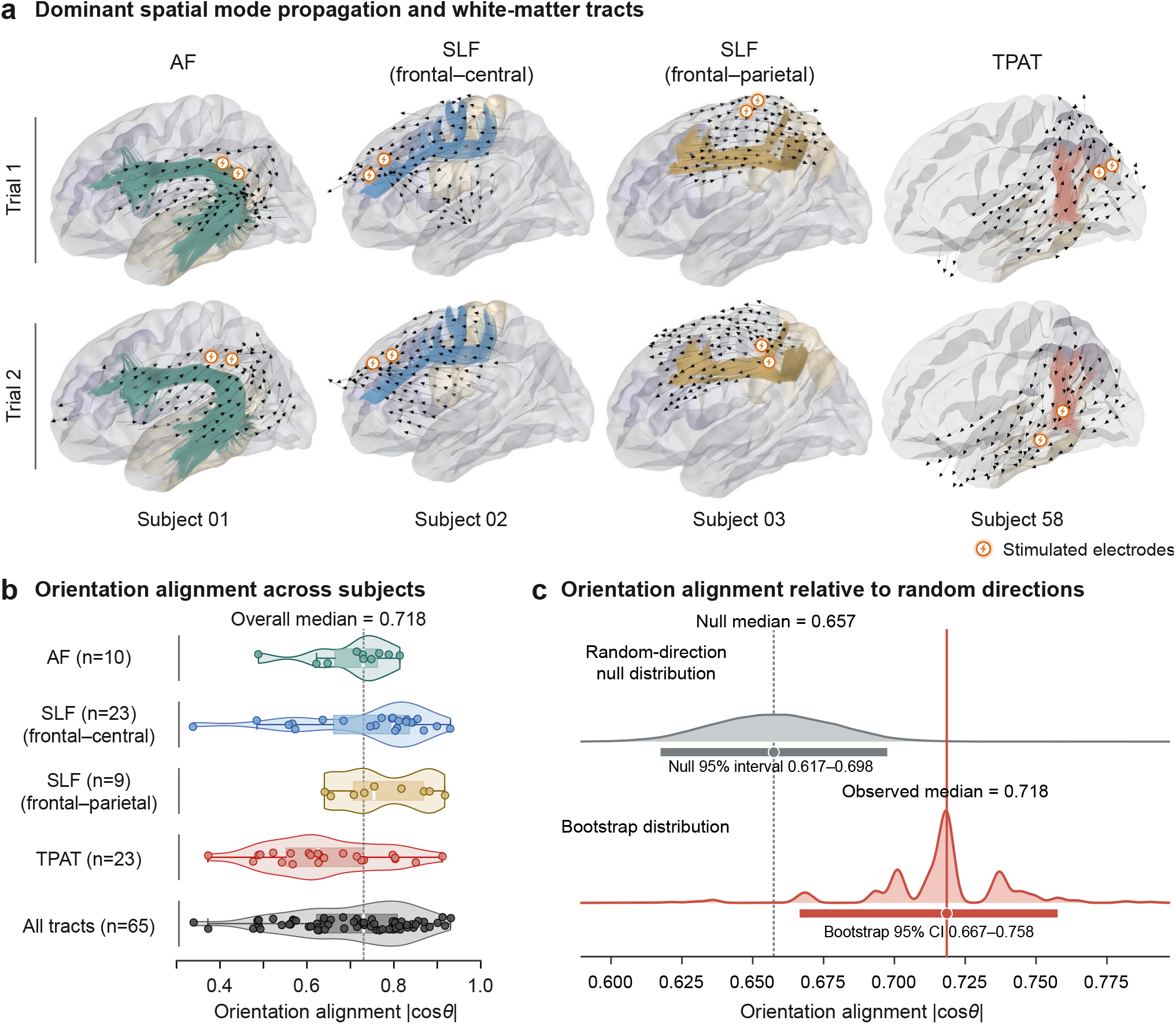
Orientation alignment between dominant cortical wave propagation and white-matter tract connection axes. **a**, Representative velocity fields of the dominant spatial mode (Mode 1) and long-range white-matter tracts. Columns show the arcuate fasciculus (AF), frontal–central and frontal–parietal segments of the superior longitudinal fasciculus (SLF), and temporo-parietal aslant tract (TPAT) for Subjects 01, 02, 03 and 58, respectively. Trial 1 (top) and Trial 2 (bottom) correspond to stimulation of different electrode pairs within each subject. Black arrows indicate dominant-mode velocity vectors, colored trajectories indicate white-matter tracts and orange lightning symbols mark stimulated electrodes. **b**, Orientation alignment between dominant-mode propagation and projected tract connection axes across subjects. Each axis links two cortical termination regions, approximating the tract’s end-to-end orientation rather than its curved trajectory. Alignment was quantified as the amplitude-weighted mean of local | cos *θ* | values, where *θ* is the angle between the locally projected connection axis and a dominant-mode velocity vector. Values of 0 and 1 indicate orthogonal and parallel or antiparallel orientations, respectively. Points represent subject-level median alignment values, and violin plots show tract-specific and pooled distributions. Sample sizes are 10, 23, 9 and 23 for AF, SLF (frontal–central), SLF (frontal–parietal) and TPAT, respectively, with 65 subjects overall. The dotted line indicates the overall median (0.718). **c**, Null distribution of the overall median alignment generated from random directions (gray, top) and bootstrap distribution of the observed overall median (red, bottom). In each of 100,000 randomizations, one three-dimensional direction was randomly assigned to each subject and projected onto the local cortical tangent planes, with the same direction used across that subject’s velocity fields. The dashed gray and solid red lines indicate the null median (0.657) and observed median (0.718), respectively. Horizontal bars show the central 95% null interval (0.617–0.698) and the 95% bootstrap confidence interval (0.667–0.758). The observed median exceeded random expectations (*P* = 0.00127, one-sided Monte Carlo test).

Each tract was represented by a tract connection axis linking its two cortical termination regions. This approximation describes the orientation of the long-range connection rather than its curved trajectory through the white matter. At each vertex, the axis was projected onto the local cortical tangent plane and compared with the dominant-mode velocity vector. The alignment score was the amplitude-weighted mean of local | cos *θ* | values, where *θ* is the angle between these vectors (Methods). Scores range from 0 for orthogonal orientations to 1 for parallel or antiparallel orientations. Scores were summarized by their median within each subject. The overall median was 0.718, with a 95% bootstrap confidence interval of 0.667–0.758 (***Figure 4***b,c).

To test whether this correspondence exceeded random expectations, we used a random-direction null model. In each of 100,000 Monte Carlo randomizations, one three-dimensional direction was assigned to each subject and shared across that subject’s velocity fields, preserving their directions, amplitudes and spatial organization. The observed median exceeded the null median (0.657) by 0.061 (*P* = 0.00127, one-sided Monte Carlo test; ***Figure 4***c). Exploratory tract-specific analyses identified correspondence above random expectations for the frontal–central and frontal–parietal SLF segments (Benjamini–Hochberg-adjusted *P* = 4.00 × 10^−5^ and *P* = 0.0334, respectively; four one-sided tests). No significant excess was detected for AF or TPAT (adjusted *P* = 0.135 and *P* = 0.937, respectively). These results support directional correspondence between the dominant spatial mode and projected tract connection axes beyond the expectations of the random-direction null model. Null-model details and group-level statistical results are provided in Supplementary Material S2.4.

### Age-related variation in cortical wave speed

We next examined the relationship between age and cortical wave speed using subject-level median estimates derived from the reconstructed velocity fields. Three of the 67 observations were excluded using the 1.5 × IQR criterion, leaving 64 subjects for regression and correlation analyses (Methods). Wave speed was positively associated with age (Pearson’s *r* = 0.549; Spearman’s *ρ* = 0.524; both *P <* 0.0001; Supplementary Material S2.2.1).

To characterize the shape of the age–speed relationship, we fitted a cubic spline model (*R*^2^ = 0.335; ***Figure 5***a). We used crossings of its first derivative at 0.002 m s^−1^ yr^−1^ and zero to define four descriptive age intervals (Methods). The fitted curve showed a relatively rapid increase over 4–15.6 years (Phase 1) and slower change over 15.6–26.9 years (Phase 2). A renewed increase over 26.9–46.8 years (Phase 3) was followed by a slight decline over 46.8–51 years (Phase 4).

**Figure 5.**
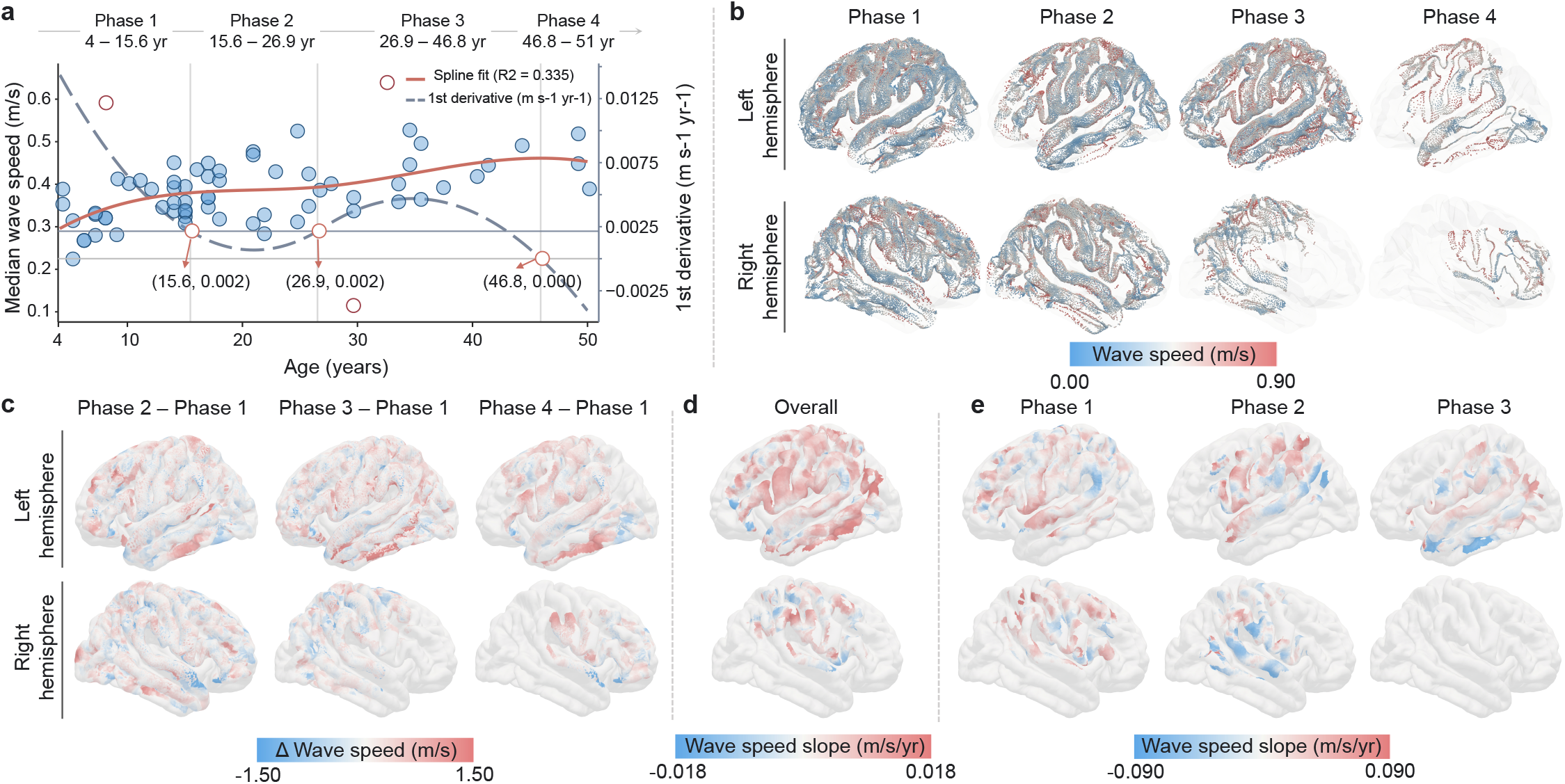
Age-related variation in cortical wave speed. **a**, Nonlinear relationship between age and subject-level median cortical wave speed. Points represent individual subjects. The solid coral curve shows a cubic spline fit (*R*^2^ = 0.335), and the dashed gray curve shows its first derivative (right axis). Dark-red open circles indicate three observations excluded using the 1.5 × IQR criterion, leaving 64 subjects for spline fitting. Open coral circles on the derivative curve mark crossings at 0.002 m s^−1^ yr^−1^ (15.6 and 26.9 years) and zero (46.8 years). These crossings define four descriptive age intervals: Phase 1, 4–15.6 years; Phase 2, 15.6–26.9 years; Phase 3, 26.9–46.8 years; and Phase 4, 46.8–51 years. **b**, Group-level spatial distributions of cortical wave speed for the four age intervals. Colors indicate wave speed in m s^−1^. **c**, Spatial differences in wave speed relative to Phase 1, calculated as each phase minus Phase 1. Red and blue indicate higher and lower wave speeds, respectively. **d**, Vertex-wise age–wave-speed regression slopes across the full cohort (4–51 years). The two maps show the lateral surfaces of the left and right hemispheres, respectively. Red and blue indicate positive and negative slopes, respectively. Only vertices meeting the predefined data-support criteria were included. **e**, Vertex-wise age–wave-speed regression slopes estimated separately within Phase 1 (4–15 years), Phase 2 (16–26 years) and Phase 3 (27–45 years). Columns correspond to the three phases, and the top and bottom rows show the lateral surfaces of the left and right hemispheres, respectively. Phase 4 was not shown because only three subjects were available. Only vertices meeting the predefined data-support criteria were included.

We then compared group-level wave-speed maps across the four spline-defined age intervals (***Figure 5***b). Relative to Phase 1, Phases 2–4 showed higher speeds in parts of the frontal, temporal and parietal cortex and lower speeds elsewhere (***Figure 5***c). To further examine spatial variation in the age–speed relationship, we performed vertex-wise linear regression. Across the full age range, regression slopes varied across cortical regions, with both positive and negative values (***Figure 5***d). Separate analyses within Phases 1–3 revealed differences in the spatial distribution of these slopes (***Figure 5***e). Phase 4 was not included because only three subjects were available. Individual wave-speed maps for subjects in the youngest and oldest intervals are provided in Supplementary Material S2.2.2. Together, these results indicate an overall positive association between age and cortical wave speed, accompanied by spatially heterogeneous age-related variation.

## Discussion

The present study introduces and applies velocity-field analysis, an interpretable framework for quantifying multiscale cortical dynamics, to reveal a hierarchical organisation of cortical waves governed by anatomical constraints, with flexible coordination potentially mediated by singular points within the default mode network. We further establish cortical wave speed as a viable biomarker of human brain development and ageing. Together, these findings may have profound implications for systems neuroscience and neural dynamical systems, as discussed below.

### Velocity fields provide finer insights into the spatiotemporal dynamics of cortical activity than conventional analyses

Velocity field analysis represents a robust and interpretable framework for characterizing cortical spatiotemporal dynamics (***Afrashteh et al., 2017; Townsend and Gong, 2018***), addressing critical limitations of conventional connectivity analyses and black-box methods (***Roberts et al., 2019; Kohoutová et al., 2020***). Compared with conventional connectivity-based approaches, which primarily capture static, low-resolution inter-regional correlations (***Reid et al., 2019; Roberts et al., 2019***), this method delivers markedly higher spatiotemporal precision, revealing transient, fine-scale neural dynamics that are often overlooked by conventional analyses (***Townsend and Gong, 2018; Muller et al., 2018; Liang et al., 2023***). Moreover, velocity fields provide a direct, physically grounded representation of cortical activity (***Afrashteh et al., 2017; Townsend and Gong, 2018; Liang et al., 2023***) by explicitly quantifying the propagation speed and direction of travelling waves, thereby linking neural signals to biologically meaningful patterns of propagation (***Liang et al., 2023; Rubino et al., 2006; Zhang et al., 2018***). It further enriches dynamic profiling by identifying subtle local patterns, including singular points and their topological properties (***Townsend and Gong, 2018; Liang et al., 2023***), which expand the repertoire of quantifiable cortical micro-dynamics beyond conventional analytical boundaries (***Roberts et al., 2019***). Importantly, velocity-field analysis does not rely on learned latent representations, which can be difficult to interpret in many machine-learning and deep-learning models (***Kohoutová et al., 2020; Tolooshams et al., 2025***). While capturing complex neural patterns, data-driven models lack mechanistic transparency and biological traceability (***Kohoutová et al., 2020***). In contrast, velocity field-derived features are fully interpretable (***Liang et al., 2023; Roberts et al., 2019***) and readily linked with cortical anatomical architectures (***Koller et al., 2024; Rubino et al., 2006***) and dynamical systems (***Liang et al., 2023; Roberts et al., 2019***). Collectively, the integration of high spatiotemporal fidelity, physical plausibility and biological interpretability renders velocity field analysis an efficient tool for dissecting multi-scale cortical dynamics, providing new insights into physiological neural computation (***Rubino et al., 2006; Zhang et al., 2018; Muller et al., 2018***) and pathological brain alterations (***Martinet et al., 2017; Smith et al., 2016; Diamond et al., 2024***).

### Hierarchical spatiotemporal organisation of cortical waves suggests rapid large-scale propagation along with local computation

Our analyses reveal a pronounced hierarchical spatiotemporal organisation of stimulus-evoked cortical travelling waves (***Figure 3***a,b), unifying large-scale propagation and fine-scale local activity to reshape our understanding of intrinsic cortical dynamics (***Liang et al., 2023; Harris and Gong, 2026; Das et al., 2026***). Through mode decomposition, we disentangle composite cortical activity into distinct dynamic components, with the dominant mode accounting for an average of 72.97% of the variance across subjects. Representative recordings exhibit broad, plane-wave-like propagation in the dominant mode, whereas secondary modes capture more heterogeneous spatial patterns, including local sources, sinks, saddles and spirals. Unlike conventional approaches that treat cortical activity as a homogeneous whole, our decomposition strategy resolves nested dynamic patterns, enabling systematic quantification of how large-scale and local neural processes coordinate across spatiotemporal scales (***Harris and Gong, 2026; Aggarwal et al., 2022***).

Remarkably, plane-wave-like propagation consistently emerges as the dominant spatial mode across different stimulation sites (***Figure 3***a,b and ***Figure 4***a), resembling the principal large-scale propagation patterns reported during spontaneous cortical activity (***Liang et al., 2023***). Despite these similarities in propagation patterns, the dominant mode accounts for a substantially greater proportion of variance in our stimulus-evoked responses than has been reported for spontaneous activity in mice (***Liang et al., 2023***). Although differences in recording modalities, cortical coverage and decomposition procedures prevent direct quantitative comparison, this contrast suggests that stimulus-evoked activity may be more strongly dominated by coherent large-scale propagation, whereas spontaneous activity exhibits more distributed contributions across dynamic modes. Together, these observations suggest that large-scale propagation represents a common feature of cortical dynamics, whose relative dominance may depend on brain state and external perturbation.

Crucially, the hierarchical wave architecture reconciles large-scale propagation with local neural computation (***Harris and Gong, 2026; Liang et al., 2023***). While dominant plane waves mediate rapid large-scale propagation to integrate widespread cortical regions (***Mohan et al., 2024; Aggarwal et al., 2022; Koller et al., 2024***), secondary modes support heterogeneous local cortical computations (***Figure 3***a,b) (***Das et al., 2026; Zabeh et al., 2023; Harris and Gong, 2026***). These local dynamics support fine-grained neural processing, including regional information encoding, local circuit modulation and context-dependent neural responses (***Zabeh et al., 2023; Harris and Gong, 2026; Das et al., 2026***). This multi-mode spatiotemporal hierarchy provides a mechanistic framework for understanding how the cortex integrates rapid inter-areal communication with regional information processing across physiological (such as alpha waves, sleep slow-waves), evoked (cortical stimulation) and pathological (epileptic seizures) conditions (***Mohan et al., 2024; Muller et al., 2018; Aggarwal et al., 2022; Diamond et al., 2024***).

### Anatomical structure guides dominant plane wave propagation, while default mode network host local singular points

Our findings uncover a structure-function mechanism governing cortical wave dynamics, whereby macroscopic plane wave propagation is constrained by anatomical architecture (***Figure 4***a–c) (***van Blooijs et al., 2023b; Koller et al., 2024***), while fine-scale local singular points are preferentially anchored within the default mode network (DMN) (***Figure 2***b,c) (***Roberts et al., 2019***). Direct cortical stimulation elicits robust travelling plane waves (***Muller et al., 2014; van Blooijs et al., 2023b***), whose spatiotemporal trajectories are strictly guided by subjacent white matter fibre tracts (***van Blooijs et al., 2023b; Koller et al., 2024***). This finding is consistent with computational models demonstrating that structural connectivity and conduction delays can generate travelling waves and shape their preferred directions (***Koller et al., 2024***). Importantly, this anatomically constrained propagation constitutes a purely physical response to external perturbation, distinct from endogenous physiological neural activity and responses from sensory stimulation (***Lyu et al., 2025***). Counterintuitively, such stereotyped plane wave patterns persist consistently across distinct stimulation sites (***Muller et al., 2014***), indicating that macroscopic cortical wave propagation is governed by invariant structural connectivity, rather than local stimulus-induced neural modulation (***Koller et al., 2024***).

Dissociation between global wave propagation and local dynamic organisation further raises a fundamental mechanistic question regarding the preferential localisation of singular points within the DMN (***Roberts et al., 2019***). As core topological landmarks of local cortical dynamics (***Roberts et al., 2019***), singular points do not distribute randomly across the cortex but selectively converge to DMN sub-regions (***Figure 2***b,c) (***Roberts et al., 2019***), implying that this network possesses unique functional properties that support dynamic neural bifurcation (***Roberts et al., 2019***) and local computational flexibility (***Xu et al., 2023; Valk et al., 2022; Fotiadis et al., 2023***). Unlike structurally constrained global plane waves, these local singular points represent physiology-driven dynamic features uncoupled from rigid anatomical propagation rules (***Valk et al., 2022; Fotiadis et al., 2023***). We propose that DMN-resident singular points serve as critical dynamic hubs for cortical functional reconfiguration (***Xu et al., 2023; Roberts et al., 2019***). These topological nodes likely operate at a subcritical energetic state, characterised by low metabolic consumption and high computational efficiency. Such sub-critical state may also enable excitatory–inhibitory (E/I) balance, endowing singular points with the capacity to mediate rapid dynamic mode switching (***Roberts et al., 2019***). While white matter anatomy establishes the rigid structural backbone for large-scale, global signal propagation (***Koller et al., 2024; van Blooijs et al., 2023b***), DMN-based singular points confer adaptive functional plasticity(***Roberts et al., 2019; Xu et al., 2023; Valk et al., 2022; Fotiadis et al., 2023***).

### Velocity fields and brain-wave speed may serve as biomarkers for brain development and ageing

As indicated by our findings, velocity fields constructed from iEEG signals may serve as a quantitative biomarker for tracking human brain development, maturation and degeneration, revealing a nonlinear age-dependent trajectory of travelling wave speed (***van Blooijs et al., 2023b***) that closely mirrors the lifelong progression of white matter development and degeneration (***Yeatman et al., 2014; Villalón-Reina et al., 2026***) (see ***Figure 5***a). Spatially resolved cortical mapping identifies distinct regional patterns of wave speed elevation and reduction across the lifespan (***Figure 5***b–e), pinpointing brain regions that undergo the most substantial micro-structural and functional reorganisation during development and ageing (***Luo et al., 2026; Lotter et al., 2024***). Such heterogeneous spatiotemporal alterations suggest that cortical dynamic remodelling follows structured region-specific trajectories, rather than uniform global change (***Luo et al., 2026; Lotter et al., 2024***), and provides fine-grained functional signatures complementary to conventional structural brain metrics.

Notably, these velocity-derived dynamic phenotypes align consistently with prior neuroimaging findings acquired through distinct modalities and analytical pipelines, cross-validating the reliability and biological validity of age-related cortical wave alterations (***van Blooijs et al., 2023b; Price et al., 2017; Yeatman et al., 2014; Villalón-Reina et al., 2026***). This methodological convergence confirms that travelling wave dynamics represent fundamental, modality-invariant neural traits tightly coupled to brain ontogeny and senescence (***Raut et al., 2021; Price et al., 2017; van Blooijs et al., 2023b***). The observed nonlinear relationship between age and wave speed further bridges macroscopic functional dynamics and micro-structural white matter changes (***van Blooijs et al., 2023b; Price et al., 2017***), capturing the transition from developmental fibre maturation and myelination refinement in early life to progressive degenerative decline in ageing brains (***Yeatman et al., 2014; Villalón-Reina et al., 2026***).

Beyond descriptive characterisation, our findings indicate that cortical travelling wave speed carries robust discriminative potential for distinguishing normative brain trajectories from pathological deviations (***van Blooijs et al., 2023b; Villalón-Reina et al., 2026***). Aberrant wave speed alterations may signal disrupted neural circuit development (***van Blooijs et al., 2023b***), impaired interregional communication (***Koller et al., 2024***), or early degenerative remodelling (***Villalón-Reina et al., 2026***), providing sensitive functional indicators of atypical development and neuro-degenerative pathology (***Villalón-Reina et al., 2026***). Building on this framework, it is feasible to construct populationbased normative models of cortical velocity dynamics across the lifespan (***van Blooijs et al., 2023b; Bethlehem et al., 2022; Villalón-Reina et al., 2026***). Such normative benchmarks would enable in-dividualised quantitative assessment (***Bethlehem et al., 2022; Villalón-Reina et al., 2026***), positioning personalised wave speed profiles as a translational biomarker for early detection, stratification and longitudinal monitoring of neuro-developmental disorders and neuro-degenerative diseases (***Villalón-Reina et al., 2026; Whitman et al., 2025***). Overall, velocity field analysis advances our mechanistic understanding of lifelong brain dynamics (***Koller et al., 2024; van Blooijs et al., 2023b***) and offers a promising and potentially clinically actionable tool for brain health evaluation.

## Methods and Materials

### Dataset description

This study utilized a publicly available cortico-cortical evoked potentials (CCEPs) ECoG dataset (***van Blooijs et al., 2023a***), comprising 74 patients (age range: 4–51 years). The dataset contains ECoG recordings acquired during single-pulse electrical stimulation (SPES). These data originate from the RESPect (Registry of Patients undergoing Epilepsy Surgery) database, collated by the University Medical Center Utrecht (UMC Utrecht), The Netherlands. Seven patients were excluded because of surface reconstruction failures or temporal segmentation anomalies, leaving 67 patients for velocity-field analyses. Two additional patients had insufficient cortical coverage for the tract connection-axis analysis, leaving 65 patients. For age-related regression and correlation analyses, three subject-level wave-speed observations were excluded using the 1.5 × IQR criterion, leaving 64 patients. Subject-specific exclusions and the age distribution are provided in Supplementary Material S2.1. Detailed dataset information and associated analysis code are publicly available on the OpenNeuro platform (accession no. ds004080).

### Data Preprocessing

A standardized preprocessing pipeline was applied to ECoG signals. Data were first band-pass filtered between 0.1 Hz and 100 Hz, followed by a band-stop filter to attenuate line-frequency interference. As indicated in the dataset metadata, channels identified as ‘bad’ were excluded, retaining only signals from ECoG electrodes. Data were re-referenced using common average reference (CAR) and baseline-corrected relative to the interval from –1 s to 0 s before stimulus onset. ECoG recordings were segmented into epochs spanning −2 to 3 s relative to stimulus onset. Epochs corresponding to the same stimulation condition were averaged to obtain CCEP responses, which were used for subsequent velocity-field reconstruction.

### Simulated data

To validate the manifold-based optical flow algorithm, we generated a synthetic dataset containing known wave patterns (i.e., singular points) following the method described in ***Townsend and Gong*** (***2018***). This approach enables the evaluation of accuracy in locating and classifying singular points. The surface was defined as an 80 × 80 grid, where *x* and *y* coordinates are uniformly distributed over the interval (−40, 40). The surface elevation *z* was modeled using a bivariate Gaussian function:

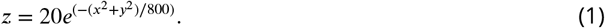

The simulated waveform *w*_sim_(*x, y, z, t*), with wave number *k* and angular frequency *ω*, was constructed as follows:

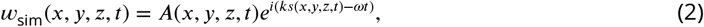

where *A*(*x, y, z, t*) governs the spatial amplitude distribution, and *s*(*x, y, z, t*) defines the spatial phase profile. By altering the functional form of *s*, different types of singular points were obtained.

Two simulation settings were considered. The first contained two moving Gaussian sources converging along diagonal trajectories, whereas the second included stationary nodes, foci and saddles (20 instances of each type). For moving patterns, the centre followed (*x*_*t*_, *y*_*t*_) = (*x*_0_ + *v*_*x*_*t, y*_0_ +*v*_*y*_*t*). Stationary patterns retained fixed centres.

For a source pattern, the spatial phase was defined as:

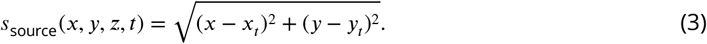

For a spiral pattern, the phase included an arctangent component:

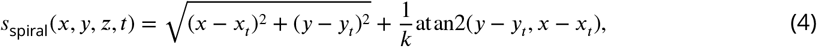

where atan2 denotes the four-quadrant inverse tangent.

For a saddle pattern, the phase was given by:

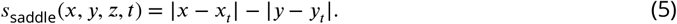

To ensure spatial localization, the amplitude *A*(*x, y, z, t*) was defined as a symmetric two-dimensional Gaussian centered at the singular point:

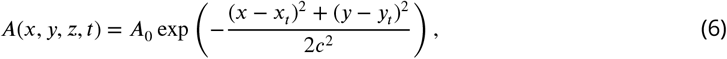

where *A*_0_ is the peak amplitude and *c* controls the Gaussian width.

In all simulations, *ω* = 2*π* × 0.01 and *k* = 2*π*∕5. Initial positions *x*_0_ and *y*_0_ were randomly selected from the domain boundary. Amplitude *A*_0_ varied between 1 and 2, and width *c* between 3 and 5. For simulations in Cortical wave singular points and their spatial association with the DMN, velocity components *v*_*x*_ and *v*_*y*_ were randomly sampled from the interval [−0.1, 0.1].

### Optical Flow Algorithm

#### Conceptual Foundations of Cortical Flow

We employed a computational framework to estimate optical flow of dynamic signals on non-flat manifolds (***Lefèvre and Baillet, 2009***). This method effectively characterizes the propagation of neural activity patterns at the centimeter scale.

Let *I*(*p, t*) denote the time-varying neural activity at point *p* on the cortical or scalp manifold *M* at time *t*. The optical flow field **V**(*p, t*) captures the local displacement over time. Under the assumption of intensity conservation (***Horn and Schunck, 1981***), **V** satisfies:

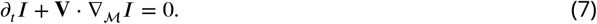

This assumption holds when neural activitiy evolves slowly relative to the high sampling rate of the EEG.

It is noteworthy that the scalar product in Equation (7) is modulated by the intrinsic geometry and curvature of *M* (***Lefèvre and Baillet, 2009***).

#### Computation

##### Theory

***Equation 7*** is an underdetermined equation (i.e., the “aperture problem”) and requires regularization. We formulated the problem as minimizing an energy functional:

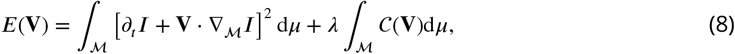

where parameter *λ* controls velocity-field smoothness. We used *λ* = 0.01 for the CCEP analyses, based on the simulation results in ***Figure 2***e.

The regularization term *C*(**V**) promotes smooth vector fields by:

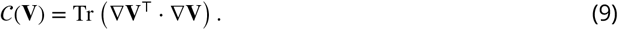

This formula extends the Horn–Schunck model (***Horn and Schunck, 1981***) to arbitrary manifolds. In the Euclidean space case (*M* = ℝ^2^), it simplifies to:

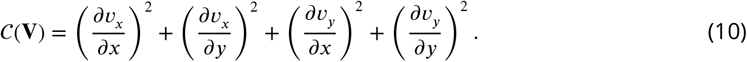

As shown in ***Schnörr (1991)***; ***Lefèvre et al. (2007)***, minimizing ***Equation 8*** reduces to solving:

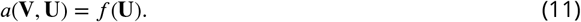

This equation applies to all test vector fields **U** in an appropriate function space, where *a* is a symmetric positive definite bilinear form and *f* is a linear form.

#### Numerical aspects

We employed the finite element method (FEM) (***Ciarlet, 2002***) to discretize this problem, following the methodology outlined in ***Lefèvre et al. (2007)***; ***Lefèvre and Baillet (2008)***. The unknown vector field **V** was represented in the basis **W**_*i*=1:*N,α*=1:2_ as tangent plane vector fields defined at mesh vertices. The coefficients *v*_*j,β*_ of **V** satisfy the following linear system of equations:

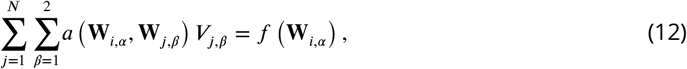

where *i* = 1 : *N* and *α* = 1 : 2. The inverse matrix of the symmetric positive definite matrix *a* **W**_*i,α*_, **W**_*j,β*_ is the optical flow estimate on *M*.

A more detailed theoretical derivation can be found in **Supplementary Material S1**. Due to the sparse distribution of intracranial ECoG electrodes, we reconstructed a continuous cortical surface model and interpolated potentials onto this surface to enable precise optical flow calculations.

### Manifold Reconstruction

A surface mesh was reconstructed based on electrode coordinates. First, a signed distance function was computed from the point cloud, and the zero-level isosurface was extracted to form an initial mesh. Subsequently, iterative Laplace smoothing was applied to refine the mesh and optimize vertex distribution. Butterfly subdivision was then employed to increase triangle density and enhance surface smoothness, ultimately yielding reconstructed manifold. Detailed experimental setup is described in **Supplementary Material 2.3.1**.

### Data Interpolation

Using radial basis function (RBF) interpolation, the potential values at the ECoG electrodes were interpolated across the entire reconstructed manifold. First, the RBF model was fitted to the scattered electrode data. Subsequently, the resulting interpolation function is evaluated at all mesh vertices, yielding a continuous potential field covering the entire manifold. Detailed experimental setup is described in **Supplementary Material 2.3.2**.

### Detection and classification of singular points

We identified singular points as locations where the reconstructed velocity field vanishes (***Effenberger and Weiskopf, 2010***). Singular points were detected on the reconstructed triangular manifold and classified according to the local velocity-field Jacobian (***Townsend and Gong, 2018***).

To detect singular points, we examined each triangular face *T* with vertices *A, B* and *C* and their corresponding velocity vectors **V**_*A*_, **V**_*B*_ and **V**_*C*_. Any point *P* within the triangle can be represented using barycentric coordinates as *P* = *αA* + *βB* + (1 − *α* − *β*)*C*. After expressing the vertex velocities in a common local tangent plane, the velocity at *P* was linearly interpolated as

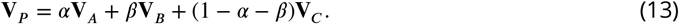

A singular point occurs when the interpolated velocity vanishes:

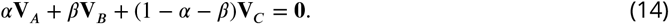

Rearranging this condition gives

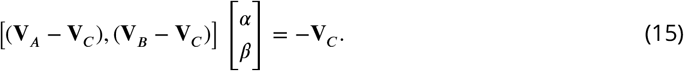

The resulting linear system was solved using least squares. A solution satisfying *α* ≥ 0, *β* ≥ 0 and *α* + *β* ≤ 1 identifies a candidate singular point within the triangle, with its location determined by the corresponding barycentric coordinates.

For each detected singular point, we estimated the Jacobian matrix of the local velocity field to characterize the surrounding flow geometry (***Ginoux, 2009; Löffelmann et al., 1996***). In local tangent-plane coordinates, the Jacobian is defined as

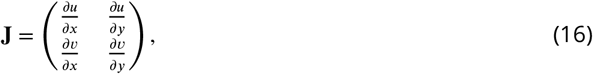

where *u* and *v* denote the two tangential velocity components. Its trace *τ* and determinant Δ are given by

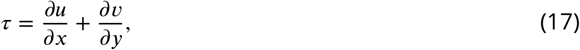

and

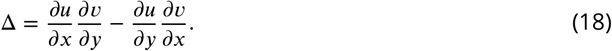

The Jacobian was estimated according to the location of each singular point. For points located at mesh vertices, neighboring velocity vectors were projected onto the local tangent plane, and directional derivatives were estimated along the tangent basis and averaged. For points within triangular faces, the vertex velocities were projected onto the corresponding face plane to estimate the local velocity gradient.

Singular points were classified according to the eigenvalue structure of **J**. Nodes satisfy Δ *>* 0 and *τ*^2^ *>* 4Δ and were classified as sources (*τ >* 0) or sinks (*τ <* 0). Foci satisfy Δ *>* 0 and *τ*^2^ *<* 4Δ and were identified as spiral patterns, whereas saddle points satisfy Δ *<* 0. These criteria distinguish the four local wave patterns examined in this study: sources, sinks, saddles and spirals.

### Velocity Field Spatiotemporal Modes Decomposition

Dimensionality reduction techniques hold significant value for identifying dominant neural dynamics (***Cunningham and Yu, 2014; Ambrogioni et al., 2017; Delis et al., 2016***). However, methods like principal component analysis (PCA) (***Abdi and Williams, 2010***) often assume spatiotemporal separability, failing to capture inseparable propagating waves (***Alexander et al., 2015***). Inspired by fluid dynamics (***Adrian et al., 2000; Taira et al., 2017***), we directly applied singular value decomposition (SVD) (***Klema and Laub, 1980***) to the velocity field to extract dominant spatiotemporal modes, as demonstrated in ***Townsend and Gong*** (***2018***).

Let *u*(*P, t*) and *v*(*P, t*) denote the velocity components in the local tangent basis at point *P* = (*x, y, z*). We reconstructed the velocity fields as the matrix **W**(*t, r*′) = [*ũ*|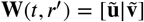], where *r*′ denotes spatial position. The singular value decomposition is expressed as:

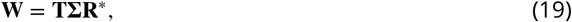

where **T** and **R**^∗^ are unitary matrices, **Σ** is a diagonal matrix with singular values *σ*_*k*_, and ∗ denotes the conjugate transpose. The *k*-th spatial mode corresponds to *k*-th and *k* + 1-th columns of **R**, explaining variance 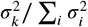. Its temporal evolution is given by the *k*-th column of **T**. To ensure interpretability, we flipped the signs of spatiotemporal mode pairs when necessary to guarantee each temporal mode has a non-negative mean.

### Directional correspondence with tract connection axes

We examined the relationship between dominant-mode velocity directions and the connection axes of four long-range white-matter pathways: the arcuate fasciculus (AF), frontal–central and frontal–parietal segments of the superior longitudinal fasciculus (SLF), and temporo-parietal aslant tract (TPAT). The analysis included 65 subjects with sufficient cortical coverage of the regions required for comparison. Each tract was represented by a connection axis linking its two cortical termination regions. This end-to-end approximation describes the orientation of the long-range connection rather than the tract’s curved trajectory.

At each vertex *i*, the three-dimensional unit connection axis **a** was projected onto the local cortical tangent plane:

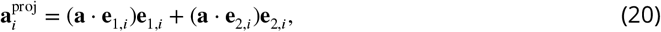

where **e**_1,*i*_ and **e**_2,*i*_ form an orthonormal tangent basis. The alignment score for each analyzed response was calculated as

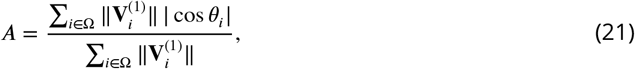

where **V**^(1)^ is the dominant-mode velocity vector, *θ*_*i*_ is its angle to the projected connection axis, and Ω denotes the included vertices. Scores range from 0 for orthogonal orientations to 1 for parallel or antiparallel orientations.

### Random-direction null model

We tested whether directional correspondence exceeded random expectations using a random-direction null model. A set of 4,096 three-dimensional directions was generated using Fibonacci hemisphere sampling. In each of 100,000 Monte Carlo randomizations, one direction was randomly assigned to each subject and shared across all trials from that subject. The assigned direction was projected onto the local cortical tangent planes, while the observed dominant-mode velocity vectors, their magnitudes and spatial organization were preserved. Alignment scores under the null model were calculated and aggregated in the same manner as the observed scores.

### Streamline Tracking of Velocity Fields on Manifolds

Streamlines are integral curves of the velocity field, tangent to the flow direction at every point (***Klausen et al., 2012; Rasmussen, 2010; Matringe and Gerritsen, 2004; Prevost et al., 2001***). They satisfy:

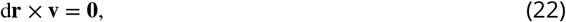

where d**r** denotes the differential arc element.

The streamline tracking algorithm proceeds as follows:

1. **Initialization:** Select seed points with non-zero velocity.
2. **Streamline tracking:**
  - For an internal point *O* on the manifold, compute the dot product of its velocity vector with the normalized vectors of neighboring points in the tangent plane. Move to the neighboring point with the largest dot product.
  - For boundary points *O*, check if velocity points outward from the mesh. If true, terminate; otherwise, proceed as above.
3. **Termination conditions:** Stop when any of the following conditions are met:
  - Velocity points toward region outside the domain,
  - Velocity is zero,
  - Closed loop detected (repeated point visits),
  - Maximum streamline length reached.
4. **Length filtering:** Discard streamlines shorter than the minimum length to ensure trajectory validity.

### Statistical Analysis of the Relationship between Age and Brain Wave Speed

We employed a multi-level statistical framework to quantify the relationship between age and cortical wave speed. After removing outliers using a combination of interquartile range (IQR) and standard deviation criteria, we analyzed data from 67 patients (age range: 4–51 years) to ensure dataset reliability.

#### Multi-Level Wave Speed Quantification

The data were hierarchically structured: for each subject (N=67), 1–8 experimental runs were acquired, with each run containing 7–728 trials. Wave speed distribution curves were constructed using the following hierarchical statistical methods:

1. **First-level analysis**: For each subject, the median wave speed at each point on the cortical manifold was calculated within a 9–50 millisecond post-stimulus time window, integrating data across all trials. This method effectively mitigates interference from outliers in local wave velocity estimates.
2. **Second-level analysis**: For each individual trial, spatial statistics were computed by taking the median wave speed across all manifold points. This reduces the spatial heterogeneity and provides a comparable, representative metric across trials.
3. **Third-level analysis**: Summarize trial-level speed within each experimental run using the average wave velocity, yielding run-specific estimates of cortical response speed.
4. **Fourth-level analysis**: Aggregate run-level estimates by subject using mean wave speed to form final subject-level metrics for age-wave speed analysis.

Age–Wave speed Statistical Modeling

We employed multiple statistical methods to describe the relationship between age and wave speed:

- **Linear regression**: Wave speed was modeled as a linear function of age to examine systematic age-related variations. Both Spearman’s (*ρ*) and Pearson’s (*r*) correlation coefficients were calculated to assess monotonic and linear relationships.
- **Spline regression**: Employ spline regression to capture potential nonlinear features in the age-wave speed relationship. Analyze first- and second-order derivatives to identify key inflection points delineating distinct developmental phases.

### Spatial analysis of cortical wave speed

Group-level wave-speed maps were constructed for the four spline-defined age intervals using the 67 subjects included in the wave-speed analysis. Subject-level wave-speed estimates on reconstructed manifolds were assigned to vertices of the common fsaverage cortical surface by nearest-neighbor mapping. At each vertex, available mapped manifold values were averaged to obtain the group estimate; vertices receiving no values remained undefined. Maps were displayed separately for the left and right hemispheres.

For spatial difference maps, each group map was extended by assigning the nearest valid source value within a 5-mm surface-geodesic distance. Equidistant nearest sources were averaged. The Phase 1 map was then subtracted from each phase map only at vertices with valid estimates in both maps. Locations outside this common support were assigned zero solely for display.

### Vertex-wise age–wave-speed regression

To characterize spatial variation in the association between age and cortical wave speed, we performed vertex-wise least-squares regression:

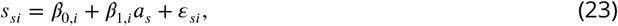

where *s*_*si*_ is the wave-speed estimate for subject *s* at vertex *i, a*_*s*_ is subject age, and *β*_1,*i*_ is the age-related slope. Within each subject, manifold values assigned to the same common-surface vertex were averaged, yielding one estimate per subject per vertex. Subject maps were extended to nearby vertices using nearest-source assignment within a 5-mm surface-geodesic distance; unsupported locations remained undefined.

Regression was performed across the full sampled age range and separately within Phases 1–3. Full-range models required at least 10 subjects and an observed age span of at least 30 years at each vertex. Within-phase models required at least five subjects and an age span covering at least half of the phase’s observed age range. Phase 4 was omitted because its three subjects could not meet the minimum sample requirement. Positive and negative slopes indicate increasing and decreasing fitted wave speed with age, respectively.

## Acknowledgments

We acknowledge the facilities and scientific and technical assistance of the Australian National Imaging Facility, a National Collaborative Research Infrastructure Strategy (NCRIS) capability, at the Swinburne Neuroimaging Facility, Swinburne University of Technology.

We thank the Big Data Computing Center of Southeast University for providing the facility on the numerical calculations in this paper.

